# The Estrogen Receptor-Related Orphan Receptors (ERRs) Regulate Autophagy through TFEB

**DOI:** 10.1101/2023.03.02.530836

**Authors:** McKenna Losby, Matthew Hayes, Aurore Valfort, John Walker, Weiyi Xu, Lilei Zhang, Cyrielle Billon, Thomas P. Burris

## Abstract

Autophagy is an essential self-degradative and recycling mechanism that maintains cellular homeostasis. Estrogen receptor-related orphan receptors (ERRs) play fundamental role in regulation of cardiac metabolism and function. Previously, we showed that ERR agonists improve cardiac function in models of heart failure and induce autophagy in cardiomyocytes. Here, we characterized a mechanism by which ERRs induce autophagy in cardiomyocytes. Transcription factor EB (TFEB) is a master regulator of the autophagy-lysosome pathway and has been shown to be important in cardiac autophagy. We discovered that TFEB is a direct ERR target gene whose expression is induced by ERR agonists. Activation of ERR results in increased TFEB expression in both neonatal rat ventricular myocytes and C_2_C_12_ myoblasts. ERR-dependent increases in TFEB expression results in increased expression of an array of TFEB target genes, which are critical for stimulation of autophagy. Pharmacologically targeting ERR is a promising potential method for treatment of many diseases where stimulation of autophagy may be therapeutic including heart failure.

## Introduction

Autophagy is a conserved degradation pathway that maintains cellular homeostasis in the cell by directing the clearance of old and abnormal proteins or organelles. A key transcriptional modulator of the autophagy-lysosome system is transcription factor EB (TFEB). TFEB is a member of the microphthalmia/transcription faction E (MiT/TFE) basic helix-loop-helix-leucine-zipper (bHLH-Zip) family that regulates transcription by binding DNA as a homodimer or heterodimer with another bHLH-Zip transcription factor.^1^ Overexpression of TFEB result in increased number of lysosomes, increased rate of degradation of autophagy substrates, induction of mitophagy and in specific disease states, halts disease progression.^1,2^ In its inactive state, TFEB is phosphorylated and localized within the cytoplasm, but upon dephosphorlation TFEB is activated and then translocates to the nucleus and regulates genes containing a CLEAR (Coordinated Lysosomal Expression and Regulation) DNA element.^1,3^ Known activators of TFEB are PPARγ coactivator 1α (PGC1α), calcineurin, and protein kinase C (PKC), while mammalian target of rapamycin complex 1 (mTORC1) and mitogen-activated protein kinase 1 (MAPK1) are known to phosphorylate TFEB, thus inactivating it.^1,4,5^ Autophagy is dysregulated in many diseases, thus activation of TFEB is believed to have potential therapeutic utility for diseases with lysosomal dysfunction and cellular clearance abnormality such as heart failure, Parkinson’s disease, Alzheimer’s disease and others.^6,7,8,9,10^ There are several compounds that have been suggested to target TFEB directly (resveratrol, curcumin analog C1, progestins) and indirectly (torin1 and rapamycin) and some of these have progressed to clinical studies.^11^ Although there are promising preliminary results with agents that activate TFEB, there are limitations with current small molecule drugs which display undesirable side effects due to broad pharmacological effects.^12^

The estrogen receptor-related orphan receptors (ERRs) are members of the nuclear receptor superfamily of ligand activated transcription factors. There are three ERR isoforms (ERRα, ERRβ, and ERRγ) that are expressed in tissues with high energy demands such as the brain, heart, muscle, and brown adipose tissue.^13^ The ERRs bind to DNA response elements, called ERREs, with a specific nucleotide sequence (TCAAGGTCA) and activate transcription of these target gene expression.^14^ ERRs regulate a number of genes involved in pathways such as fatty acid oxidation (FAO), TCA cycle, oxidative phosphorylation (OXPHOS), and mitochondrial biogenesis.^15,16^ The ERRs are orphan receptors, with no known natural ligands, however, they have a high level of basal activity in the absence of any ligands.

Although ERR inverse agonists have been known for some time, it has been challenging to develop high affinity ERR agonists that can be used as chemical tools to target the ERRs, particularly for in vivo studies. ERRα and ERRγ inverse agonists (such as XCT790 and tamoxifen) were among the first chemical tools targeting ERRs to be identified, but both of these have issues with specificity.^17,18,19^ GSK4716 was the first ERRβ/γ agonist described but its potency is relatively weak and it has limited utility in vivo.^20,21^ We recently described two novel high affinity ERR agonists (SLU-PP-332 and SLU-PP-915) that have sufficient pharmacokinetic properties to use in animal models of disease.^22,23^ SLU-PP-332 has an EC50 of 98 nM with ERRα, 230 nM ERRβ, and 430 nM ERRγ providing for a degree of alpha selective, while SLU-PP-915 has EC_50_ values of approximately 400 nM for each.

Recently, we found that treatment of mice with heart failure with these ERR agonists improved cardiac function and noted a substantial increase in expression of genes in the autophagy pathway.^23^ Furthermore, we observed that SLU-PP-915 increases autophagic flux in neonatal rat ventricular myocytes (NRVMs).^23^ Here, we examined the role of ERRs and pharmacological activation of ERRs in regulation of autophagy and found that TFEB, a critical master regulator of autophagy, is a direct ERR target gene and mediates ERR induction of the autophagy pathway.

## Results

### ERR agonists induce the expression of autophagy genes

In our previous study in mouse models of heart failure, we noted that treatment with either ERR agonists, SLU-PP-332 and SLU-PP-915, resulted in increased expression of an array of genes within the autophagy pathway.^23^ One of these genes, *Tfeb*, plays a master regulatorly role in this pathway as a transcription factor (TFEB) that induces the expression of multiple genes in the autophagy pathway. In order to begin to examine the mechanism by which ERRs regulated TFEB expression we treated neonatal rat ventricular myocytes (NRVMs) with SLU-PP-915 and assessed *Tfeb* expression by qPCR. As shown in Figure 1A, SLU-PP-915 treatment (2.5 μM, 72 h) significantly increased *Tfeb* expression in NRVMs by ~40% relative to DMSO treated cells. TFEB protein expression was also significantly increased by ~50% with SLU-PP-915 (2.5 μM, 72h) treatment (Figure 1B-C). We also assessed this effect in an additional cell type, the mouse C_2_C_12_ myoblast cell line. TFEB gene and protein expression was also increased significantly after treatment with SLU-PP-915 (5 μM, 3 h, 72 h) to a similar degree as we had observed in NRVMs (Figure 1D-F). We sought to compare the level of induction of TFEB by an ERR agonist to a physiological inducer of autophagy. A key physiological inducer of autophagy is nutrient deprivation and we compared 72 h treatment of SLU-PP-915 with an acute starvation (48 h treatment with DMEM, 24 h low glucose media) and found SLU-PP-915 induces TFEB expression to a similar extent as starvation (Figure 1G-H).

**Figure 1:**
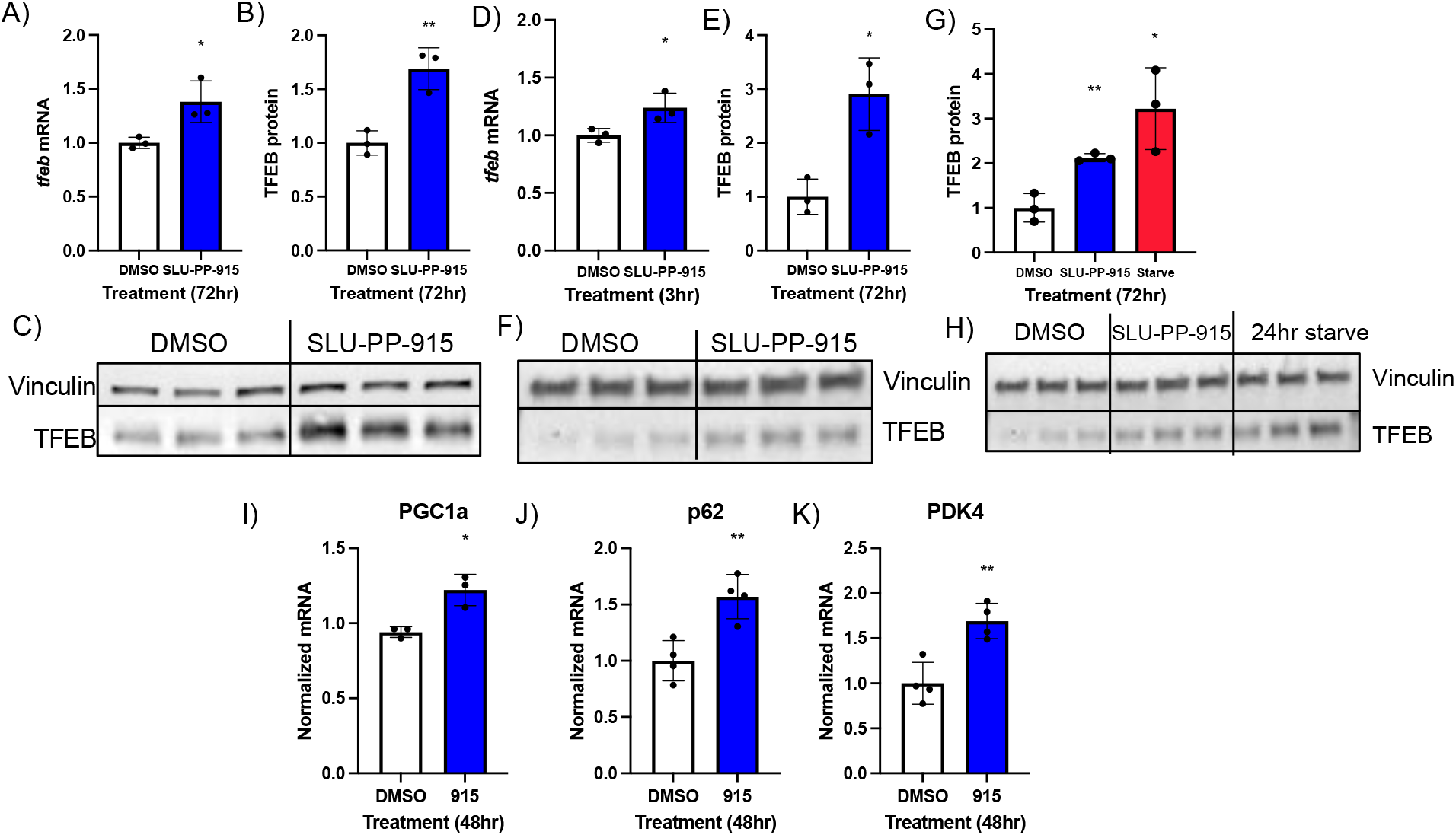
ERR agonists induce the expression of autophagy genes and proteins. A) Gene expression of TFEB increases after 72 h treatment with SLU-PP-915 (2.5 μM) in NRVMs. B) Protein expression of TFEB increases after 72 h treatment with SLU-PP-915 (2.5 μM) in NRVMs. C) Representative immunoblotting of TFEB in NRVMs quantified in B). D) Gene expression of TFEB increases after 3hr treatment with SLU-PP-915 (5 μM) in C_2_C_12_ cells. E) Protein expression of TFEB increases after 72 h treatment with SLU-PP-915 (5 μM) in C_2_C_12_ cells. F) Representative immunoblotting of TFEB protein in C_2_C_12_ cells quantified in E. G-H) TFEB protein expression was compared with SLU-PP-915 treatment (5 μM, 72 h) and starvation (24 h low glucose media). I-K) ERR target genes are significantly increased after 72 h treatment with SLU-PP-915 (5 μM) in C_2_C_12_ cells.

Other ERR target genes and autophagy genes, *Ppargc1a, p62*, and *pyruvate dehydrogenase kinase 4 (Pdk4*), were found to be significantly increased after treatment with SLU-PP-915 (5 μM, 48 h) in C_2_C_12_ cells (Figure 1 I-K). We also utilized a qPCR autophagy array to assess the broad effects of pharmacological activation of ERR on autophagy genes in NRVMs and as illustrated in Supplemental Fig. 1, SLU-PP-915 treatment (5 μM, 72 h) induced the expression of genes involved in autophagosome formation, cargo sequestering, autophagosome closure, and fusion with the lysosome.

Given the constitutive transcriptional activation activity of ERRs, we would also expect that they may play a role in maintaining basal expression of *TFEB*. The HEK293 cell line expresses predominately ERRα (proteinatlas.org) and we generated an ERRα KO CRISPR HEK293 cell line and observed that *TFEB* expression was significantly lower in the KO line vs. the WT line consistent with ERRα driving basal *TFEB* expression (Supplemental Fig. 2).

**Figure 2:**
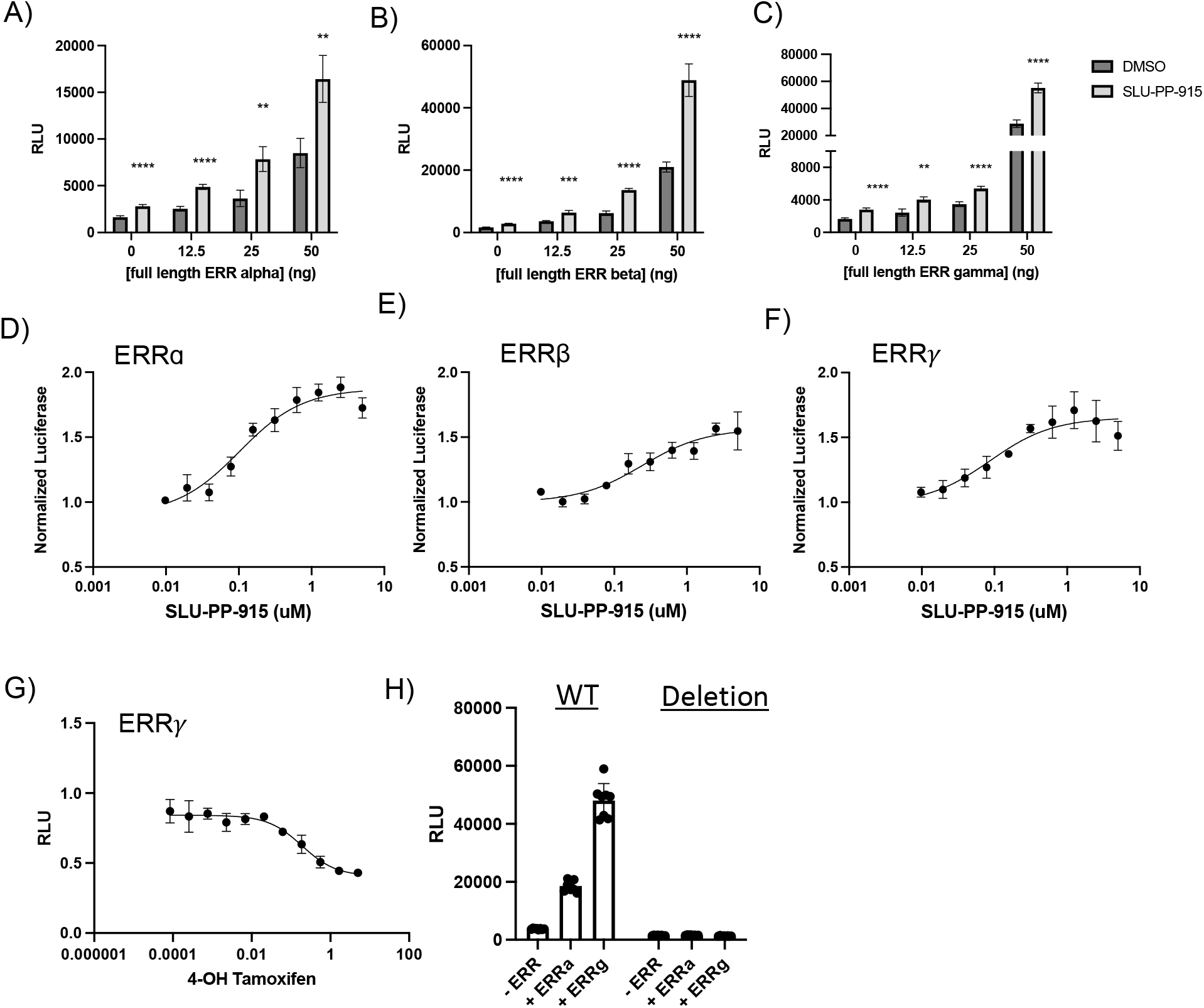
ERR regulates the transcription of TFEB through binding to ERRE’s in the TFEB promoter. A-C) Luciferase co-transfection assay in HEK293 cells show dose response dependency of luciferase signal by varying the amount of full length ERR receptor, with or without treatment of SLU-PP-915. D-F) Luciferase co-transfection assay in HEK293 cells show all three isoforms of ERR display binding to TFEB promoter in a dose response manner with SLU-PP-915. G) ERRγ inverse agonist 4-hydroxy tamoxifen with TFEB-luc decreases luciferase signal in a dose response manner. H) When the ERREs are deleted from the TFEB promoter luciferase signal is decreased.

### ERR response elements are located in the promoter of the TFEB gene

Provided that pharmacological activation of ERR results in increased TFEB expression and TFEB is an upstream regulator of many of the other genes induced by the ERR agonists, we first examined if TFEB might be a direct ERR target gene. Examination of ChIP-seq data for ERRγ (WT and CRISPR ERRγ KO hiPSC-CMs (GEO Series accession number GSE113784))^24^ we observed multiple ERRγ binding signals in the *Tfeb* promoter that are present in the WT cells but absent in the ERRγ KO cells (Supplemental Fig. 3). The sequence corresponding to the ChIP-seq ERRγ peaks were subjected motif analysis software, FIMO and MCAST, that identified two ERRE’s in the TFEB promoter (Supplemental Fig. 4A).^25,26^ Upon cloning this promoter sequence into a pGL4.23 luciferase containing plasmid (named *tfeb*::*luc*), we performed a luciferase co-transfection assay with full length ERRα, β, and γ and observed that each of the ERRs was able to increase the basal reporter gene driven by the *TFEB* promoter in a dose-depenent manner consistent with the observation of a functional ERR binding site (ERRE(s) located within the promoter (Figure 2 A-C)). Importantly, SLU-PP-915 activated *TFEB*-promoter driven luciferase signal in a dose response manner for each of the full length ERRs (Figure 2D-F). Of note, we observe a significant increase in luciferase signal without addition of drug, suggesting basal activation of TFEB by ERR. We also observed that the ERR agonist derived from a distinct chemical scaffold, SLU-PP-332, induces a similar response (Supplemental Fig. 5). An additional pharmacological approach was taken to confirm ERRs transcriptional regulation of TFEB by examining the effects of an inverse agonist on TFEB-luc activity. 4-hydroxy tamoxifen, an ERRγ inverse agonist, suppressed transcription driven by the TFEB reporter consistent with ERR mediated regulation of TFEB expression (Figure 2G).

We then deleted a region (1600pb) of TFEB promoter containing two of the putative ERREs in order to assess the ERR/ERRE-dependence of the reporter gene expression. (Supplemental Fig. 4B). Without this region containing the putatives ERREs present in the *tfeb::luc* plasmid (named *tfeb-deletion::luc*), we observed significant decrease in gene expression compared to the WT construct consistent with the constitutively active ERR driving basal *Tfeb* expression (Figure 2H). Moreover, we did not observe any increase in luciferase activity with overexpression of either ERRα or ERRγ (Figure 2H) suggesting a key role of this region in ERR-dependent TFEB activation.

### ERR agonist increases total cellular TFEB

In addition to assessing TFEB expression via western blot, we used immunofluorescence (IF) allowing for both quantitation and localization of the protein. Specifically, we assessed TFEB expression by IF in C_2_C_12_ cells treated with SLU-PP-915 (5 μM, 48 h). We found that SLU-PP-915 increased total TFEB by approximately 2-fold (Figure 3A-E). Upon comparison of TFEB in the nucleus between control and SLU-PP-915 treated C_2_C_12_ cells, we found a significant increase in nuclear TFEB (Figure 3F). Given that TFEB is a transcription factor increased TFEB in the nuclear would be expected to drive increased activity.

**Figure 3:**
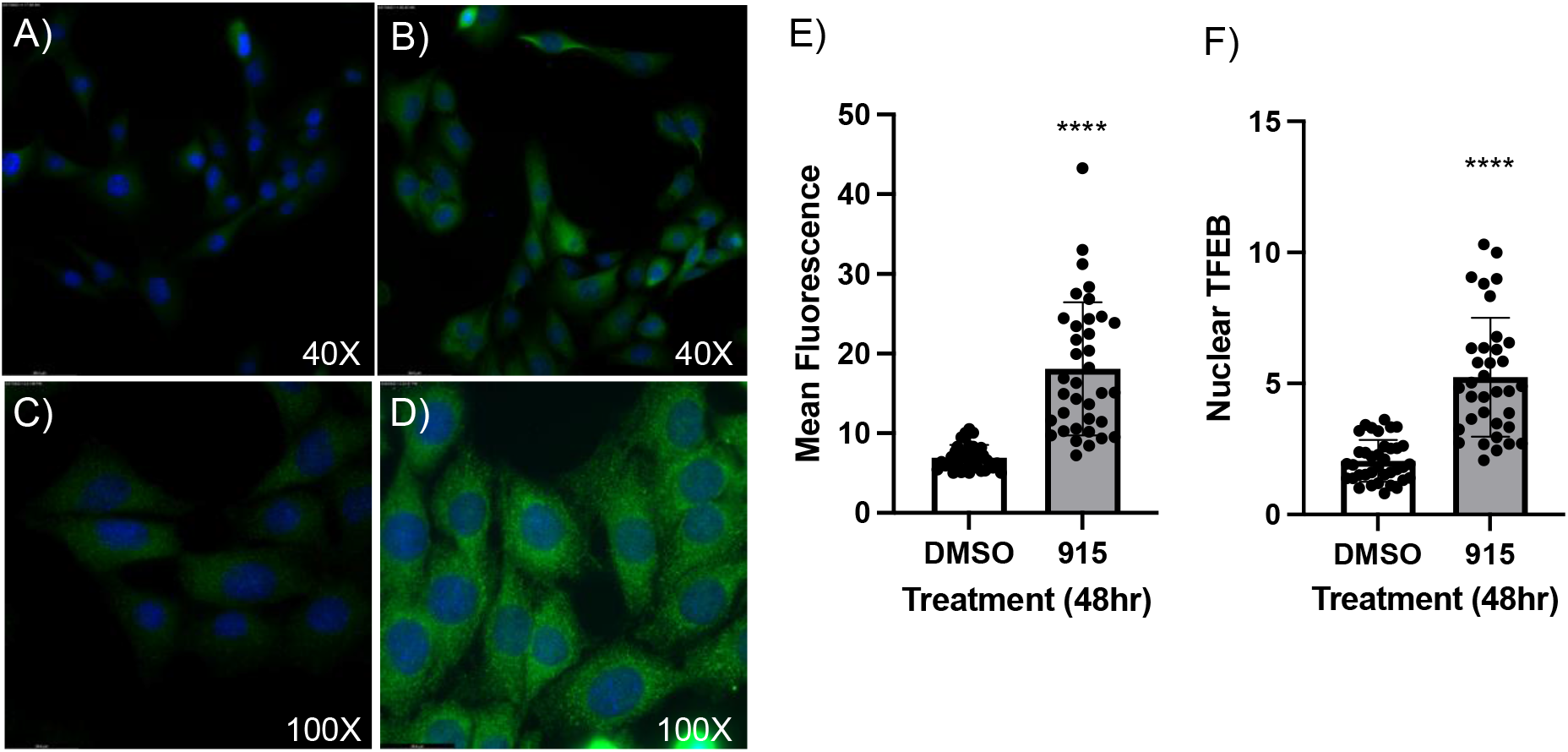
Total mean fluorescence of TFEB increases upon treatment with SLU-PP-915. A-B) Immunofluorescence staining of TFEB in C_2_C_12_ cells treated for 48 h with DMSO or SLU-PP-915 (5 μM) at 40x. C-D) Immunofluorescence staining of TFEB in C_2_C_12_ cells treated for 48 h with DMSO or SLU-PP-915 (5 μM) at 100x. E) Mean fluorescence of the whole cell is quantified utilizing ImageJ software. F) The fluorescence of nuclear TFEB is quantified utilizing ImageJ software.

### SLU-PP-915 treatment induces expression of TFEB target genes

Given that TFEB plays a critical role in regulation of an array of genes that modulate the autophagy/lysosome pathway, we compared known TFEB target genes to differentially expressed genes identified in NRVMs treated with SLU-PP-915 (5 μM, 72 h) by RNA-seq.^23^ Activation of ERR by SLU-PP-915 induced a large array of TFEB target genes in pathways such as autophagy/lysosome, transport, and metabolism (Figure 4 A-C). Several TFEB target genes that were identified to increase expression in response to SLU-PP-915 by RNA-seq in NRVMs were validated by qPCR and found to significantly increase in expression with SLU-PP-915 treatment in C_2_C_12_ cells as well (Figure 4 D-G). Lysosomal-associated membrane protein 1 (*Lamp1*) was one of these genes identified in the comparision and we examined LAMP1 in greater detail and confirmed induction of LAMP1 protein expression by SLU-PP-915 (Figure 4 H-I).

**Figure 4:**
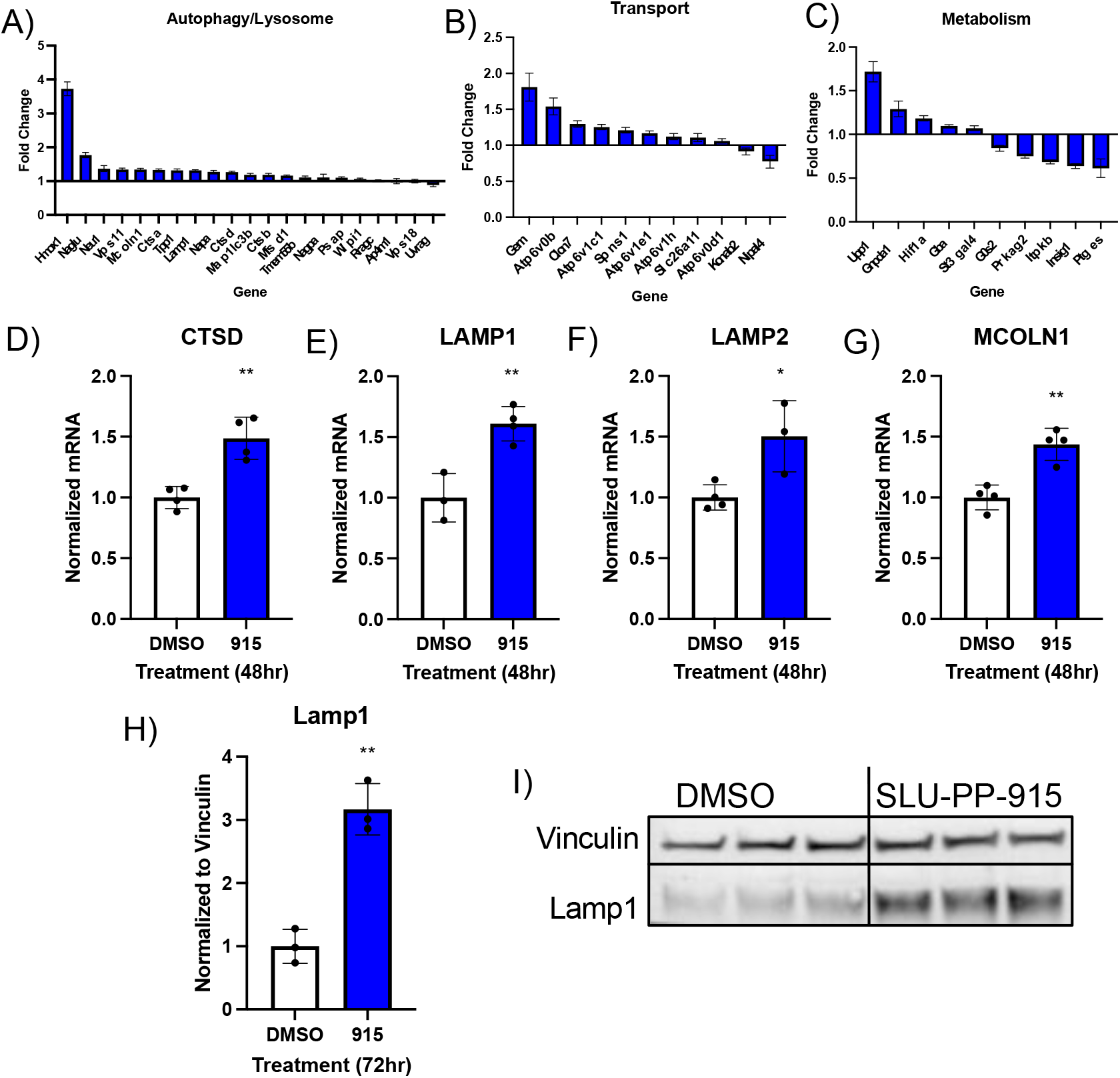
TFEB target genes are significantly increased upon treatment of ERR agonist SLU-PP-915. A-C) TFEB target genes are modulated in SLU-PP-915 treated NRVMs. The fold change is shown as SLU-PP-915 CPM/ DMSO CPM. D-G) SLU-PP-915 (5 μM) impact on TFEB target genes were validated with qPCR in C_2_C_12_ cells treated for 48 h. H) Protein expression of Lamp1, TFEB target gene. increases after 72 h treatment with SLU-PP-915 (5 μM) in C_2_C_12_ cells. I) Representative immunoblotting of Lamp1 in C_2_C_12_ cells.

## Discussion

We previously observed that the ERR agonist SLU-PP-915 increases transcription of autophagy related genes and increases autophagic flux.^23^ Based on our analysis of the autophagic genes induced by activation of ERR, we hypothesized that TFEB induced by pharmacological activation of the ERRs may be the key factor underlying induction of the pathway. Here, utilizing both primary cardiomyocytes (NRVMs) and mouse myoblasts (C_2_C_12_) we demonstrated: 1) Agonizing ERR with SLU-PP-915 increases gene and protein expression of TFEB, to a physiological level of autophagy induction as compared with starvation; 2) ERR binds to the *Tfeb* promoter and the *Tfeb* promoter confers ERR responsiveness to a luciferase reporter gene; 3) when the region containing 2 putative ERREs is deleted from the *Tfeb* promoter, the ERR responsiveness of the reporter is eliminated; 4) after treatment of ERR agonist, TFEB target genes are induced, suggesting a signaling cascade event.

Genetic loss of function in TFEB KO mice is embryonic lethal demonstrating the vital importance of this transcription factor.^27^ Autophagy flux is reduced in both tissue specific TFEB KO in the muscle of mice and by siRNA transfection of cardiomyocytes *in vitro* illustrating the importance of TFEB in autophagy.^28,29^ Furthermore, overexpression of TFEB both enhances autophagy flux and lysosomal biogenesis and restores autophagy flux in knockout models.^29,30,31^ Increased TFEB activity in cardiomyocytes may be therapeutic in the treatment of heart failure given the decrease in autophagy in this disease, but this approach should include caution given possible side effects that may result from excessive TFEB activity. Some studies show that cardiac specific overexpression of TFEB results in detrimental effects such as increased markers of pathological hypertrophy and premature death with evidence of heart failure due to decreased mitochondrial bioenergetics, increased calcium regulation and pro-fibrotic pathways.^32,33^ Although overexpression studies have their advantages, their clinical relevance may not be realistic given that the overexpression of TFEB may provide activity much higher than could be obtaining in a therapeutic situation. In our pharmacologyical/therapeutic case, increasing expression and transcriptional activity of TFEB would not be at the levels of genetic overexpression, thus may not cause the deleterious effects observed in that model.

Our observation of TFEB target genes modulated by SLU-PP-915 treatment suggests a signaling cascade where pharmacological activation ERR leads to increased transcription of TFEB. Increased TFEB expression then leads to enhanced expression of TFEB target genes and thus increased autophagy. We propose a possible feedback mechanism that may occur between TFEB and ERR (Figure 5). Our data suggests that ERR regulates transcription of TFEB, however, a recent study highlights that TFEB directly regulates ERRα in the context of endometrial cancer.^34^ Thus, both ERR and TFEB may have the capacity to regulate one anothers activity allowing for optimal control of autophagy depending on the cellular environment.

**Figure 5:**
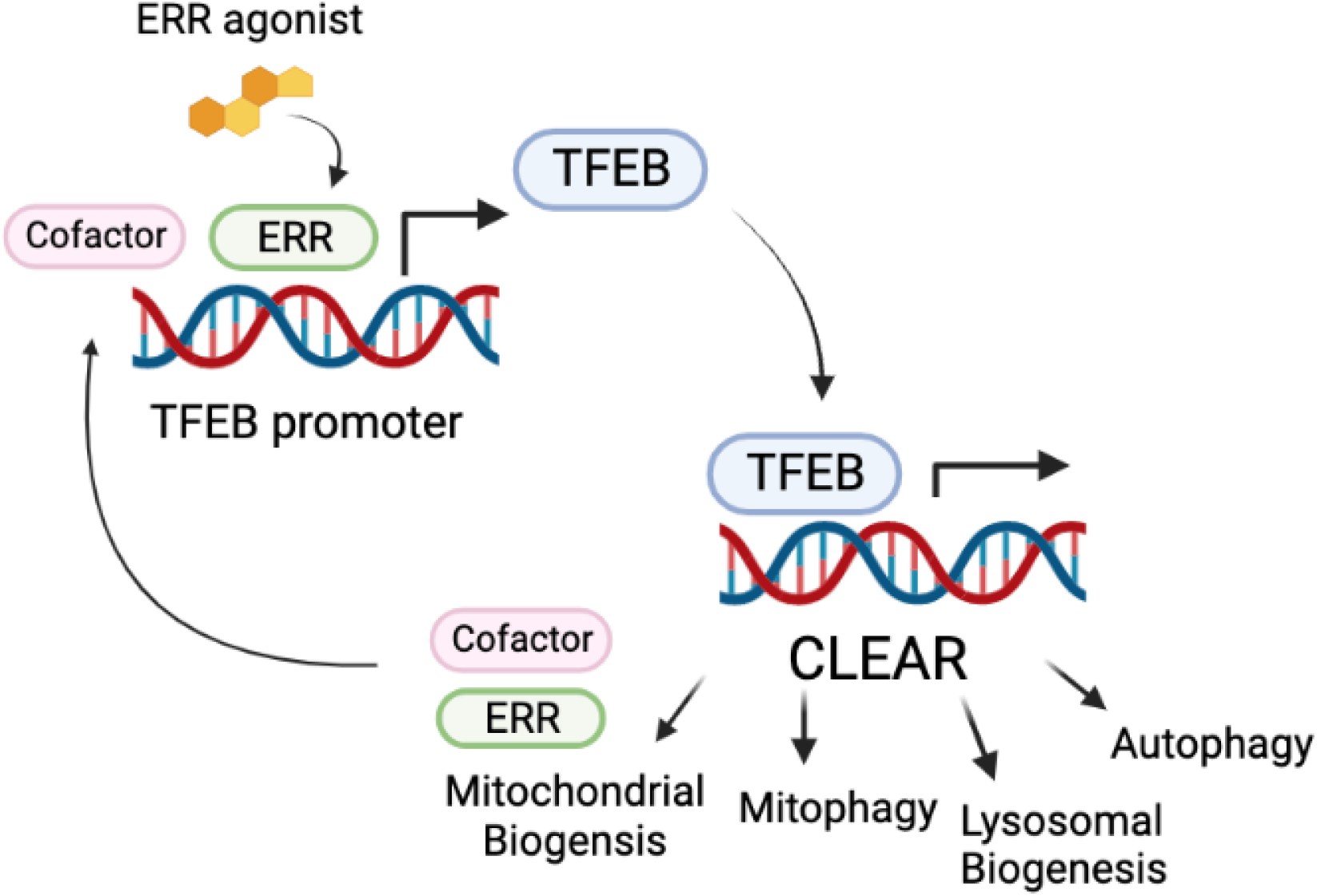
Proposed mechanism for ERR-TFEB regulation. Activating ERR with ERR agonist increases expression of TFEB through binding an ERRE in the TFEB promoter. TFEB regulates CLEAR gene network though binding its response elements. TFEB regulates gene pathways including autophagy, lysosomal biogenesis, mitophagy, and mitochondrial biogenesis. ERRs are transcribed by TFEB, thus completing the loop.

Our results suggest that ERR agonists may have therapeutic utility in treatment of heart failure since a pharmacologic agent that increases cardiac autophagy may ameliorate the disease by decreasing mitochondrial damage, oxidative stress, inflammation, and apoptosis.^35^ According to the CDC, 659,000 people die from heart disease each year in the United States, making it the leading cause of death.^36,37^ In a TAC (transverse aortic constriction) model of heart failure, autophagy was decreased suggesting that an increase in autophagy may be cardioprotective.^38^ Moreover, SLU-PP-915 has been shown to improve heart function in TAC model.^23^ There are no current treatments for heart failure that target cardiac autophagy and targeting ERRs may be a mechanism to target this pathway in treatment of heart failure. TFEB’s role in cardiac autophagy is a topic of interest due to the findings that TFEB is decreased in patients with heart failure, protects against proteotoxicity, normalizes desmin location, reduce inflammatory cytokines, diminishes myocardial dysfunction post injury, and more.^39,40,41,42^ ERR may be an ideal therapeutic target for treatment of heart failure because not only does it target autophagy-lysosome pathway as described in this work, but also targets the metabolic dysfunction that occurs in the failing heart. Genetic studies of cardiac specific ERRα/γ deletion in mice display heart failure phenotype with dysregulated metabolic, contractile, and conduction fucntion.^43^ In heart failure there is a shift in fuel utilization to glycolytic metabolism from oxidative metabolism, so targeting ERR with a small molecule agonist will increase the oxidative metabolism machinery and reverse the metabolic pathways inhibited in the failing heart. By targeting both of these deficits in heart disease, it may be possible to target multiple cellular abnormalities associated with heart failure simultaneously.

## Experimental Procedures

### NRVM isolation

All animal work was performed in accordance with protocols approved by Washington University in St Louis Animal Care and Use Committee (IACUC). NRVMs were isolated and cultured as previously described.^23^ The hearts from D0-3 rat pups were isolated and rinsed in HBSS (ThermoFisher Scientific, 14025092) before placed in trypsin-EDTA 0.05% (ThermoFisher Scientific, 25300120) at 4°C overnight. The following morning the cardiomyocytes undergo digestions with collagenase type II (ThermoFisher Scientific, 17101015) in HBSS, followed by a 90 min pre-plating incubation at 37°C with 5% CO_2_. The supernatant was collected, and the cells seeded in Gibco DMEM (ThermoFisher Scientific, 10566024) with 10% FBS, 1% pen/strep, and 100uM BRDU (Sigma-Aldrich, B5002). After 48hrs the media is changed to DMEM with 0.1% ITS (PeproTech, 00102) and the cardiomyocytes are used for experimentation.

### Luciferase Co-transfection Assays

HEK293 cells were cultured in Gibco DMEM (ThermoFisher Scientific, 10566024) supplemented with 10% FBS at 37°C with 5% CO_2_. Cells were plated in 96-well plates at a density of 2×10^4^ in 50 μL media 24 hours pre-transfection. The cells were transfected using Lipofectamine 2000 with TFEB-luc and FL-ERR plasmids (Fisher Scientific, 31985088). 24 hours post-transfection the cells were treated with ERR agonist or DMSO control. After 24 hours, luminescence activity was detectected using Promega OneGlo Luciferase Reagent (Promega). Luminescence values were normalized to DMSO control and data was analyzed using GraphPad Prism.

### Western blotting

The compounds used for NRVM treatment were prepared at 10 mM in DMSO, which was then diluted to 10 μM, 5 μM, or 2.5 μM for *in vitro* assays. Stock solutions were aliquoted and stored at −20 °C. NRVMs were lysed with RIPA buffer (VWR, 97063-270) with protease inhibitor cocktail (Sigma-Aldrich, 05056489001) and protein concentration determined with Pierce BCA assay kit (ThermoFisher Scientific, 23227). Samples were mixed with Tru-Page loading buffer (Sigma-Aldrich, PCG3009) and DTT before boiling for 5 minutes at 95 °C. Samples were loaded on 4-20% gels with TGS buffer (Fisher Scientific BP1341-1). The protein was transferred to PVDF membrane (Bio-Rad, 1704156EDU). Membranes were incubated with primary antibodies overnight at 4°C (Supplemental Table 1). Secondary antibodies were diluted 1:3000 and incubated for 1 h at room temperature.

### Nuclear Fractionation

C_2_C_12_ cells were treated for 72 h with SLU-PP-915 at 2.5 μM and Torin at 250nM for 1 h, then nuclei and cytoplasmic fractions separated using CelLytic^™^ NuCLEAR^™^ Extraction Kit (Sigma-Aldrich, NXTRACT-1KT). The resulting protein was run on a western blot and normalized to loading controls Histone H3 for nuclear fraction and alpha-tubulin for cytosolic (Sup. Table 1). The ratio was taken of nuclear to cytosolic TFEB.

### RNA and DNA extraction

For RT-PCR from NRVM cell culture, RNA was extracted using RNeasy Mini Kit (Qiagen, 74106) per manufacturers protocol. DNA from NRVMs was extracted using QIAmp DNA mini kit (Qiagen, 51304) per manufacturers protocol.

### RT-PCR

Reverse transcriptase was performed using iScript Reverse Transcription Supermix (Quanta, 95047). Sybr Green Supermix (Thermo Fisher, 4309155) was used was RT-PCR and analyzed in Quant Studio 5 Real-Time PCR System. Expression levels were normalized by PPIB (rat) and 36B4 (mouse) control and ΔΔCt-method for calculation. Primer sequences are included in Supplemental Table 2.

### Immunofluorescence

C_2_C_12_ cells were plated on coverslips in 12-well plates at 10k cells/well. After 24 h the cells were treated with SLU-PP-915 at 5 μM for 48 h. The cells were fixed with 4% paraformaldehyde (PFA) for 15 minutes then rinsed 3x with PBS. The cells were permeabilized with 0.1% Triton X-100 for 15 minutes, then rinsed 3x with PBS. The cells were blocked with 3% bovine serum albumin (BSA) in PBS for 1 h at room temperature. The cells were incubated overnight at 4°C with TFEB antibody (1:200) (Fischer Scientific, 13372-1-AP). The following day the primary antibody was removed from the cells and rinsed with PBS 3x. Secondary antibody (1:1000) (Fisher Scientific, 111-545-144) was incubated at room temperature for 1 h and the cells were rinsed 3x with PBS. The cells were incubated for 10 minutes with DAPI (1 μg/mL), then rinsed 3x with PBS. The coverslips were mounted with Prolong Gold Antifade (Invitrogen, P36934) overnight at room temperature. Slides were imaged with Leica Fluorescent Microscope and analyzed using imageJ software.

### Motif analysis

TFEB was visualized on Chromosome 6 of GRCh38 using the Interactive Genomics Viewer. ERRγ Chip-seq BigWig tracks from WT hiPSC-CMs were displayed along the genome.^24^ The sequence of a region (chr6:41735945-41733302) of occupancy within the TFEB promotor was exported to the fasta format. A motif search was performed on the sequence using Find Individual

Motif Occurrences (FIMO) and Motif Cluster Alignment and Search Tool (MCAST) from MEME Suite (v5.4.1).^25,26^ The ERRE sequence, TCAAGGTCA, was queried allowing for variation in the first and second base.

### Statistical analyses

All data are presented as mean with standard deviation of the mean with statistical significance defined as p< 0.05. t-tests were performed in GraphPad Prism.

## Abbreviations

BAF: bafilomycin A1
ChIP-seq: chromatin immunoprecipitation with high-throughput sequencing
CLEAR: Coordinated Lysosomal Expression and Regulation
DMEM: Dulbecco’s modified Eagle’s medium
DMSO: dimethyl sulfoxide
ERR: estrogen related receptor
ERRE: estrogen related receptor response element
FAO: fatty acid oxidation
GFP: green fluorescent protein
hiPSC-CM: human induced pluripotent stem cell-derived cardiomyocytes
MAPK1: Mitogen-activated protein kinase 1
MOI: Multiplicity of infection
mTORC1: mammalian target of rapamycin complex 1
NCOR1: Nuclear Receptor Corepressor 1
OXPHOS: oxidative phosphorylation
PGC1a: peroxisome proliferator-activated receptor gamma coactivator 1-alpha
PKC: protein kinase c
RFP: red fluorescent protein
TAC: transverse aortic constriction
TCA: tricarboxylic acid cycle
TFEB: transcription factor EB

## Acknowledgements

We thank Genome Technology Access enter (GTAC) in the Deparment of Genertics at Washington University School of Medicine for performing the RNA-sequencing. This work is supported by Washington University Center for Autophagy Therapeutics and Research (WUCATR).

## Notes

### Competing Interest Statement

The authors have declared no competing interest.

