## Supplemental Data for "The Estrogen Receptor-Related Orphan Receptors (ERRs) Regulate Autophagy through TFEB"

| Antibody | Source |
|---|---|
| TFEB (Western Blot) | Cell Signaling Technology, 83010s |
| TFEB (IF) | Fischer Scientific, 13372-1-AP |
| Lamp1 | Abcam, ab208943 |
| Vinculin | Cell Signaling Technology, 18799S |
| Alpha Tubulin | Abcam, ab15246 |
| Histone H3 | Cell Signaling Technology, D2B12 |
| Goat anti-Rabbit IgG, Alexa Fluor 488 | Fisher Scientific, 111-545-144 |

Supplemental Table 1: Antibodies used in western blotting and immunofluorescence assays.

| Gene (mouse (m) or rat (r)) | F Primer | R Primer |
|---|---|---|
| r TFEB | 5’-CGGTCACTGAAGGACAGAGT-3’ | 5’-GGAGTCTTTAAGCGTGGGCT-3’ |
| r PPIB | 5’-ACCAATGGCTCCCAGTTCTT-3’ | 5’-CTCCACCTTCCGTACCACAT-3’ |
| m TFEB | 5’-CCACCCCAGCCATCAACAC-3’ | 5’-CAGACAGATACTCCCGAACCTT-3’ |
| m PGC1a | 5’-CGGAAATCATATCCAACCAG-3’ | 5’-TGAGGACCGCTAGCAAGTTTG-3’ |
| m p62 | 5’-CCTCTGAGTCTCGGGAATTTCA-3’ | 5’-GACTTACTGCACGTTTGGGC-3’ |
| m PDK4 | 5’-CCGCTGTCCATGAAGCA-3’ | 5’-GCAGAAAAGCAAAGGACGTT-3’ |
| m Ctsd | 5’-GCTTCCGGTCTTTGACAACCT-3’ | 5’-CACCAAGCATTAGTTCTCCTCC-3’ |
| m Lamp1 | 5’-CAGCACTCTTTGAGGTGAAAAAC-3’ | 5’-CCATTCGCAGTCTCGTAGGTG-3’ |
| m Lamp2 | 5’-TGTATTTGGCTAATGGCTCAGC-3’ | 5’-TATGGGCACAAGGAAGTTGTC-3’ |
| m Mcoln1 | 5’-CTGACCCCCAATCCTGGGTAT-3’ | 5’-GGCCCGGAACTTGTCACAT-3’ |
| m 36B4 | 5’-ACCTCCTTCTTCCAGGCTT-3’ | 5’-CCCACCTTGTCTCCAGTCTTT-3’ |

Supplemental Table 2: Primers used in gene expression studies.

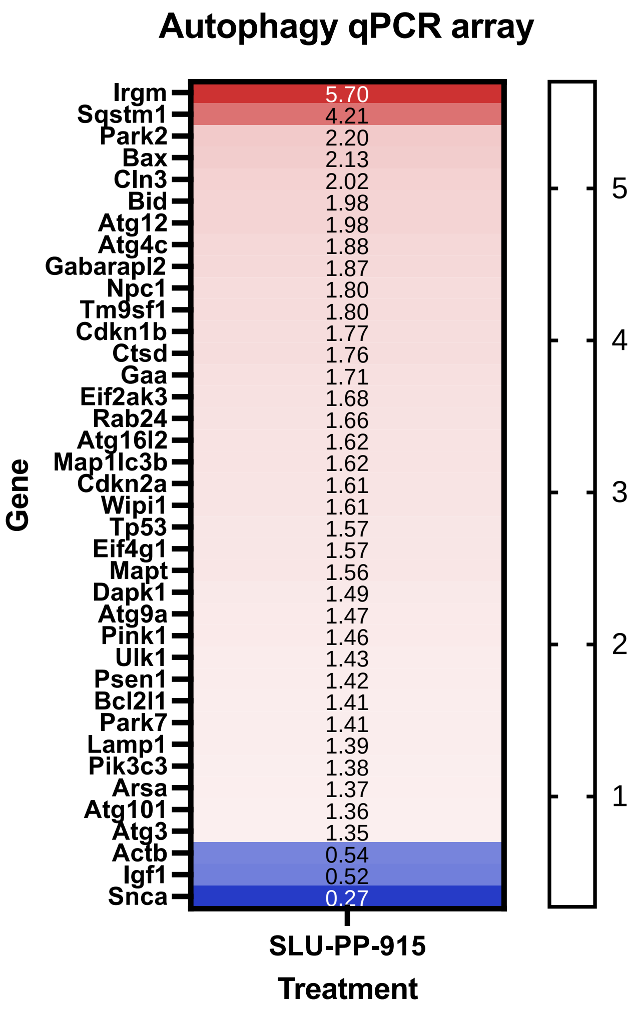


Supplemental Figure 1: Genes involved with autophagy pathway are differentially expressed in NRVMs treated with ERR agonist. NRVM were treated for 72 h with SLU-PP-915 (5 μM) , then ran on an autophagy qPCR array plate.

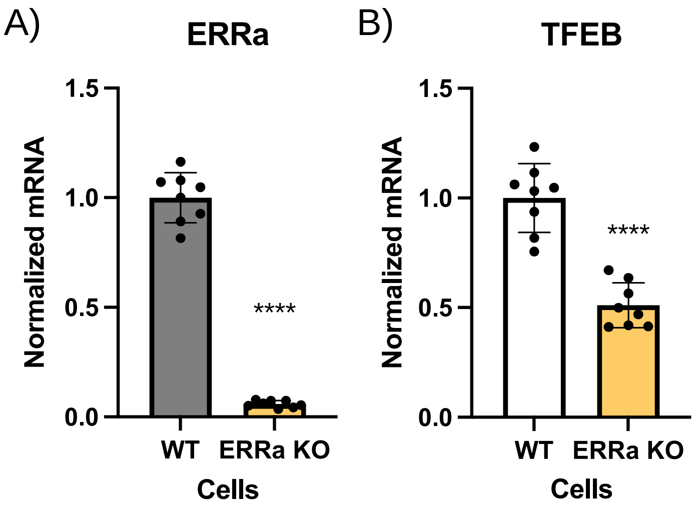


Supplemental Figure 2: TFEB expression is decreased in HEK293 cells without ERR a. A-B) qPCR characterization of CRISPR KO ERR a. C) TFEB gene expression is significantly decreased in ERR a CRISPR KO cells.

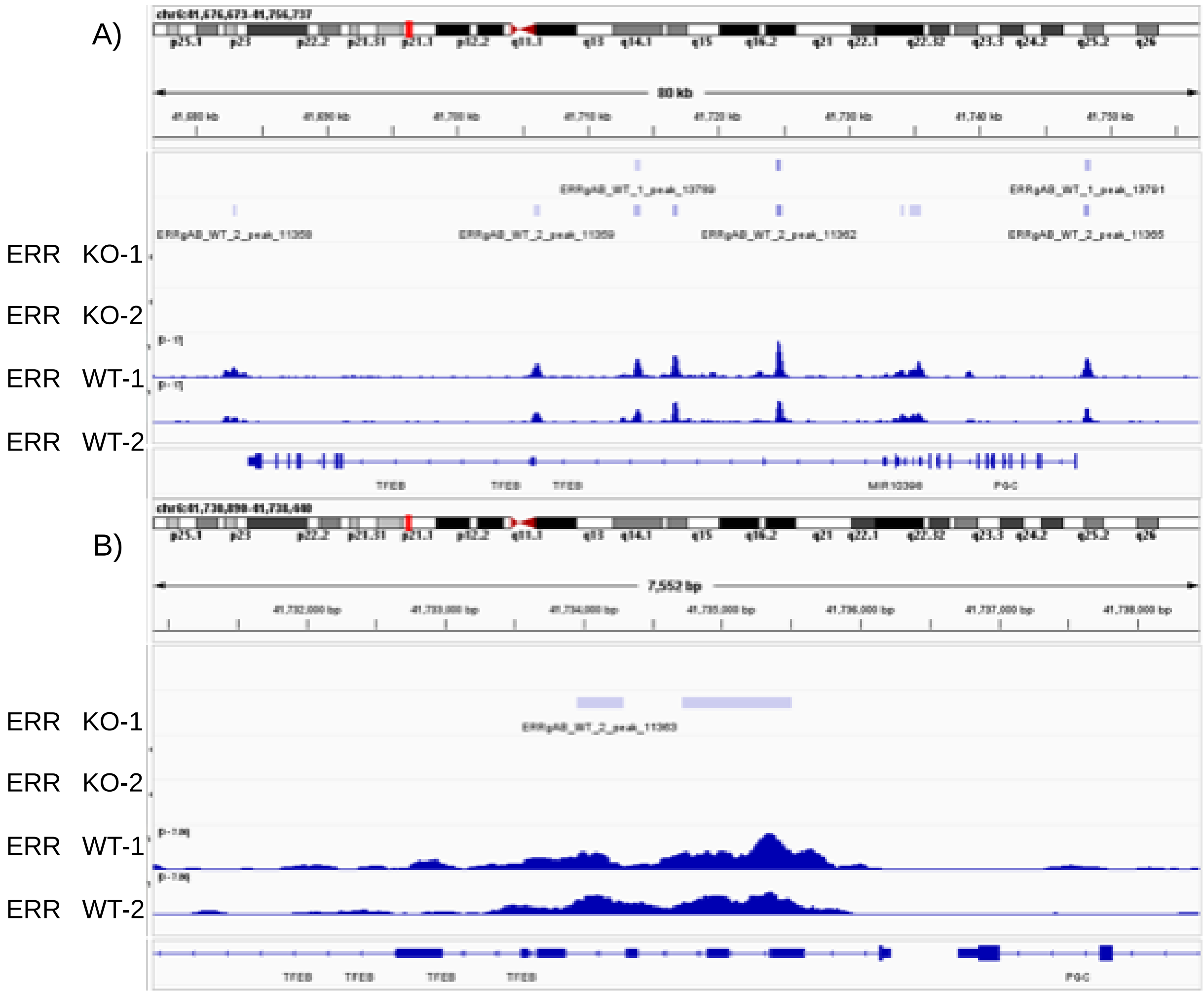


Supplemental Figure 3: ERR binds to the TFEB promoter as seen by ChIP-seq.[24] A) ChIP-seq of ERR $\gamma$ from hiPSC-CMs reveal multiple peaks associated with the TFEB gene. B) Zooming into the promoter region shows the peaks containing ERRE sites.

A)

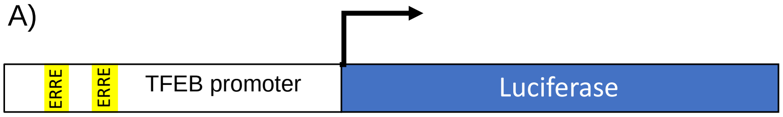


ACACCATCTGGGACCCTCTCTTTCTTAGACTGCCGAAGGACAGCAGGCTGCAGGTTCCTTTTAGGGGGTCGGACAGGCTACAGAATGTATCTTAGTGAGCCCTGGTTT
GGGACTTTACTGAGAGGAGAAAGAAAACTTGCAAAGAGGTAGAAGTGGATCAGAGGTTCCAAGTCAAGAGCCTCTTTCCCTGTAAGCCTTAGTCCGCTGAGCGGGA
CTACAAATCCCGGCGTGCCTAGCGGGACTGCAGCGAGAACTACGCTTCCCAGTGGGCGCCGAGGCAGAGCGGCTGGGAGGCGGGGCGGGTGCTGCCCTCTGCCGG
GCGCGGAGGGAAGTGTGCGTGGAGTGACAGCCGGAGCCCGGGCGCCGGGCGCGGGGCTCGGTGACAGCGGAGGCGGCGGCCCGGGCGGGACGCAGGGAGCGG
CCGCGGCGCCGACGGCGGCGGGAAGGGCGGACGGGGCGGGCCTCCCTGGTGGCACGGGGCCGGAGCAGGCCGGGAGCAGGTGGCGGGCACAGCTTGCGGGCCA
GGCACCCGAACTTGCGACAAGTTGCCGGAGCCGCGGGGCGCGGGCGGCGGACAGATTGACCTTCAGAGCGAGGTGAGCGGGACCGGGGAGCGAGAGGCACGAG
GAGGGCCCGGGACCCCCGACCCAGGCTCCCTGGCCCCTATCCCGTGGGTCCCGGAGACAGGTGGGAGGGTGGGGGATGCCGGCAGGAGAGGGAAGCAGGACCGT
GCGCGCCGCCACCCACCCGCAGTCCGTCCGCCCCCGGGCGCCCAGCGCGTGGCCCGAGCGGCGCTGCCCGCCGAGCCGGGAGCCCCCTCGGCCCCTCCCTTCAGGCG
CGGGCCGGGGGCGGCTTCTCCGCGACCTTATGTAACCGGGCGGGAGGGGCCGGGCGGGCATGGGCCTTCCCGGCCCGGAGCTGGGAGTCGAAGGGGCGGGAGGC
GTGATGGTGAACTCGCAAGAAGTTTGAGGGACGCGCGGGCCCCGCGCCCACTCCCCCTCCACCGGACACGGCTGGGGCCGGCGATGCCTGAGAGGGGGTCGGAGG
TAGGGGAGGTGTGCGGGGGCTGAGTCTGGCCAGGCCAGCGGGGGGAGTGGGAGGGCCAGAGCGGCGCCACGCCCCGGGGGCTGGGCTGATGGGGGGGCGGGT
GGAAGTACCATCTCTCCCTCCCTTCCCGTCCCTTCCTCCAGCTCGGTCTCTTGAAGAAGCCGCCCCTGGCCCCGTCCCCTCGAGTCCCTGGGCAGTCTCTTTCTCCCCGT
GGGTTATTCTATGACCCTGGGGAACACCAACCCCGGAGCGCTCCCTCCGTGCGCCCCGTTGTCTGCCTCCCCCCGGGTTCCCAGCTGCCTGCCCGCCCCCCGAGTTTCC
CCCTCCCCCCGTCTCGGTCCCGAATCCCGGAAAACACGCCCCCTTCCCCCCGATCCCCCGAGTCCGTGCCGCCCACCCCTGGGCTGTCCCGACCAGCTCTGGCCCAGCG
CGTCCGCTCGCCTCCTGGAAGGGAGCGGGCAGATGCCAGCTCGGCCCCCTCCCCGCGCGGGGTTCCGGCTTCACGCACGGCTGCCCCGAGCTGCTCCGGGCTGCGCG
AGCAGGTTCGCGCCGCAGCCGCCTGGCGCGCCCCAGCCCCGCGCCCCGGGTGAGGAAGGGAGGGAGCCCGGGGCGGGGCTTGTCTGCGCGAACCCCGCTGGGCCC
AGGCCCCCGCCCTCCGCTCTCCGCCAGCCCCTGGCCGCTGCGCCTTCCCCCGAACCCCCCTCAGCTCCAGCCGTCGTTCTCTCGCCTGTCTGTTCTGGGGACGGTTCAG
CGCGGAGCTTCTGCACTCAACTTGAGCGCAGTTTTTGGCTTAGAGCTCGCTGGGGAAGTGCAGGTTAATTACTGCCCAGCTCCCATCCTCTGCCCTCCCCCCATCCCGA
GTGAGGGGCTGCTTCATCCCTCAGGGAAAGGGACAAGACGGATGCAGGCCCCCAGTGACACCTCTGGGGTGTGGCCTCAGGCCCCTGCACCCGCCCTGGCCGAGAT
TCCATTGGCCCAGACCTGGCACTGTTGGGCAGCTGGGGATGAGGTTGGAGGCTGGTCTCTGGTTTCCTTCTGGGTGGACAGATGCGGATGCGCTCGGTTAGAACGCA
GTTCTTGGTGCCACGACTGGGAGACAGTCCTCAGGCCCTCCCCTCAGGGAGCTCACTGTCCGGGGAGGGACAGCCAGCAATAGTTCCTGGCCGGCCAGGAGAGGCA
GAGGCAGAGGAAGGGAAGGGTCAGCTGCCTTGAGGTCCCTGCCTGCTAACCTGACGGTATGCTTTCACCATGCCGGTGACCCCAGCCCCAGGACCACGAGCAACTC
GCCTGGCTCCCTTCTCTCCGTCTGCCTCCTGGCCGCGGGGCCCGGAGAGCTGGGAGCCAGAGGGTGTAAATGAAGGGCAGAGCTGTTTGGGTGACTCCATGCTCTGC
TTCTCCCCGAGGGAAGCCAATGGTGCTTACCATTTACAAGGCTTGTTACAGCTGCAGGGTTATTTTGTCCTCAACTCAGCTCTGGGAGACGGAGCCAAATGGGGACCC
AGGTATCTTGGTTCCGGGTGCCCTGGAAGGAGTTCAGGGGATAGGGCAGGGGACACCCTAGGTGGGCACTGAATCTAGGGAGAGCCAACTGGGCTTT

B)

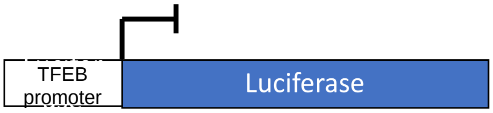


CCAGCCCCGCGCCCCGGGTGAGGAAGGGAGGGAGCCCGGGGCGGGGCTTGTCTGCGCGAACCCCGCTGGGCC
CAGGCCCCCGCCCTCCGCTCTCCGCCAGCCCCTGGCCGCTGCGCCTTCCCCCGAACCCCCCTCAGCTCCAGCCGTC
GTTCTCTCGCCTGTCTGTTCTGGGGACGGTTCAGCGCGGAGCTTCTGCACTCAACTTGAGCGCAGTTTTTGGCTTA
GAGCTCGCTGGGGAAGTGCAGGTTAATTACTGCCCAGCTCCCATCCTCTGCCCTCCCCCCATCCCGAGTGAGGGG
CTGCTTCATCCCTCAGGGAAAGGGACAAGACGGATGCAGGCCCCCAGTGACACCTCTGGGGTGTGGCCTCAGGC
CCCTGCACCCGCCCTGGCCGAGATTCCATTGGCCCAGACCTGGCACTGTTGGGCAGCTGGGGATGAGGTTGGAG
GCTGGTCTCTGGTTTCCTTCTGGGTGGACAGATGCGGATGCGCTCGGTTAGAACGCAGTTCTTGGTGCCACGACT
GGGAGACAGTCCTCAGGCCCTCCCCTCAGGGAGCTCACTGTCCGGGGAGGGACAGCCAGCAATAGTTCCTGGCC
GGCCAGGAGAGGCAGAGGCAGAGGAAGGGAAGGGTCAGCTGCCTTGAGGTCCCTGCCTGCTAACCTGACGGT
ATGCTTTCACCATGCCGGTGACCCCAGCCCCAGGACCACGAGCAACTCGCCTGGCTCCCTTCTCTCCGTCTGCCTC
CTGGCCGCGGGGCCCGGAGAGCTGGGAGCCAGAGGGTGTAAATGAAGGGCAGAGCTGTTTGGGTGACTCCAT
GCTCTGCTTCTCCCCGAGGGAAGCCAATGGTGCTTACCATTTACAAGGCTTGTTACAGCTGCAGGGTTATTTTGTC
CTCAACTCAGCTCTGGGAGACGGAGCCAAATGGGGACCCAGGTATCTTGGTTCCGGGTGCCCTGGAAGGAGTTC
AGGGGATAGGGCAGGGGACACCCTAGGTGGGCACTGAATCTAGGGAGAGCCAACTGGGCTTT

Supplemental Figure 4: A) The sequence of the TFEB promoter containing the two ERREs (highlighted in yellow) was cloned into a luciferase containing vector and named “TFEB:luc”. B) The ERREs were deleted from the TFEB promoter, along with the sequence between them, and cloned into a luciferase containing vector named “TFEB:deletion”.

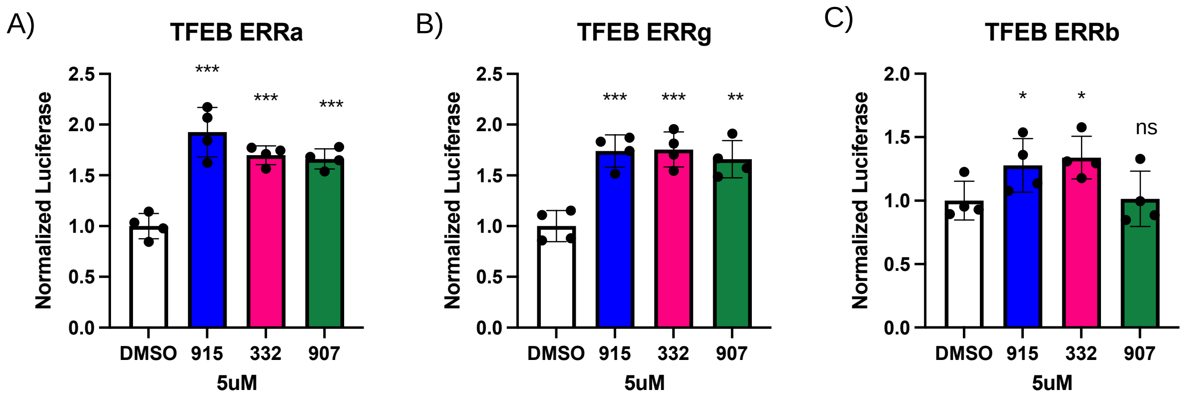


Supplemental Figure 5: Luciferase signal is significantly increased for three ERR agonists with different scaffolds. Co-transfection in HEK293 cells with FL A) ERR $\alpha$, B) ERR $\gamma$, or C) ERR $\beta$, and TFEB-luc, treated at at 5 $\mu$M for each agonist significantly increases luciferase signal, showing specificity for ERR in this reporter system.
